# Platinum Cross-linked Collagen Matrices with Tunable Stiffness as a Platform to Investigate Cellular Mechanosensing

**DOI:** 10.64898/2026.04.21.720034

**Authors:** Shinichiro F. Ichise, Yuki Taga, Kazumasa Fujita, Takaki Koide

## Abstract

The mechanical properties of the cellular microenvironment are key regulators of cellular physiology. Although the field of cancer mechanobiology has attracted attention, the availability of matrix systems with independently and precisely tunable mechanical properties remains limited. Our group previously developed a collagen gel with enhanced mechanical strength by cross-linking collagen molecules using a platinum complex. In this study, we investigated the tunability of the mechanical properties of the platinum cross-linked collagen gel (PCG) and demonstrated that mechanical parameters can be controlled by varying the amount of the platinum complex. In addition, we examined how matrix mechanical properties modulate the phenotypes of lung adenocarcinoma A549 cells using the collagen matrix. Although A549 cells exhibited significant morphological alterations on stiffer matrices, these changes were not accompanied by classical epithelial-to-mesenchymal transition (EMT). Instead, they were associated with the upregulation of diverse gene expression related to cancer malignancy. We focused on maternal embryonic leucine zipper kinase (MELK) whose gene expression increased on stiffer matrices. Consistently, A549 cells cultured on stiffer matrices displayed enhanced sensitivity to a MELK-targeting anticancer drug. These findings highlight the potential of the matrices with tunable mechanical parameters not only to provide variety of physiological microenvironment but also to advance anticancer drug screening when combined with gene expression analysis.

**Graphical abstract:** 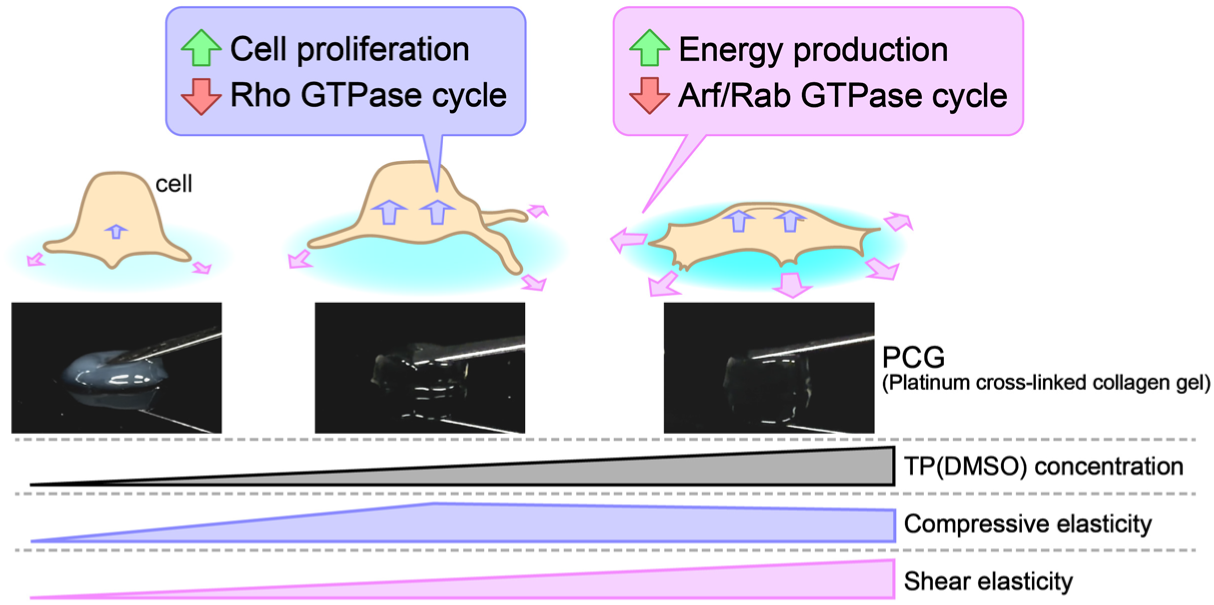

**Highlights:**

- Platinum cross-linked collagen gel enables independent tuning of compressive and shear elasticities.
- Cellular functions may be regulated by matrix mechanical parameters through distinct mechanisms.
- Correlation analysis between matrix mechanical parameters and cancer cell gene expression provides a rational strategy for therapeutic drug screening.

## 1. Introduction

Cells sense their surrounding environment and regulate various physiological functions. Such regulation has traditionally been understood through receptor-mediated signal transduction triggered by extracellular matrix (ECM) components, soluble factors, or neighboring cell membrane proteins. Recently, accumulating evidence has shown that cellular functions are also regulated by mechanical cues from the surrounding matrix [1,2]. Consequently, studies investigating the relationships between matrix mechanical properties and cellular phenotypes, particularly in cancer cells, have attracted attention and contributed to the establishment of the field of cancer mechanobiology.

The stiffness of the peritumoral matrix is not only clinically associated with poor prognosis[3,4] but also has been reported to be a factor driving malignant phenotypes in *in vitro* models and in transplantation studies using mice [5–7]. Integrins are major cell-surface receptors that sense the mechanical environment, and the downstream signaling and following nuclear translocation of Yes-associated protein (YAP) are known as a representative mechanotransduction pathway [8,9]. In cancer cells, activation of integrin–YAP signaling in response to a stiff matrix has been reported to induce epithelial-to-mesenchymal transition (EMT), a key mechanism underlying cancer invasion and metastasis [10,11]. EMT is a canonical process in cancer progression, through which epithelial cancer cells lose cell-cell adhesion, acquire mesenchymal-like phenotypes, and exhibit enhanced migratory capacity. EMT is classically induced by stimulation with transforming growth factor-β1 (TGF-β1), and is accompanied by characteristic molecular changes, including cadherin switching from E-cadherin to N-cadherin, upregulation of vimentin, and increased expression of EMT-related transcription factors such as SNAIL, SLUG, TWIST1, and ZEB1/2 [12,13].

Collagen is a major component of the ECM that not only contributes to the physical properties of the matrix but also is known to regulate malignant phenotypes of cancer cells through collagen receptor-mediated signaling [14,15]. Type I collagen, which can be abundantly extracted from skin and tendons, has been used as a scaffold for cell culture. Due to its gel-forming properties, it has also been widely employed as a 3D matrix that provides a physiological mechanical environment. Shintani *et al*. reported fundamental insights into the effects of collagen on cancer phenotypes and their mechanisms [16–18]. Notably, they demonstrated that collagen stimulation induces upregulation of endogenous TGF-β3 expression in lung adenocarcinoma A549 cells, thereby triggering EMT through an autocrine activation.

In addition, Fujisaki *et al*. reported that the fibrillar structure of collagen plays a critical role in EMT induction in A549 cells [19]. In the study, fibrillar collagen was shown to induce cadherin switching more efficiently than non-fibrillar monomeric collagen. However, they have also shown that collagen stimulation suppresses the expression of vimentin, leading to the conclusion that fibrillar collagen induces a partial EMT. Nevertheless, it should be noted that the mechanical contributions of the collagen matrix used in that study were not addressed.

Our group previously reported a collagen gel material with high transparency and improved physical strength through cross-linking of collagen molecules by a platinum complex [20]. Cisplatin and transplatin, anti-cancer platinum drugs, transform into a dimethyl sulfoxide (DMSO)-complex upon dissolution in DMSO. These complexes are capable of coordinating with the side chains of methionine and histidine residues on a collagen triple helix, thereby cross-linking and polymerizing collagen molecules [20,21]. The platinum complex and the resulting gel have minimal cytotoxicity. Therefore, the gel could be utilized as a cell culture scaffold. In this study, we aimed to test the hypothesis that the mechanical parameters of the gel could be tuned by the amount of the platinum complex. Furthermore, we investigated how the mechanical parameters regulate EMT or related cellular behaviors in A549 cells.

## 2. Materials and methods

### 2.1. Materials and cell culture

Porcine skin-derived type I atelocollagen acid solutions (3 mg/mL or 5 mg/mL) were obtained from Koken (Tokyo, Japan). Bovine skin-derived type I atelocollagen was procured from Nippi (Tokyo, Japan). Sequencing grade-modified trypsin was purchased from Promega (Madison, WI, USA). A transplatin-derived dimethyl sulfoxide (DMSO) complex, referred to as TP(DMSO), solution was prepared by dissolving transplatin (BLD Pharmatech, Shanghai, China) in DMSO (Fujifilm Wako Pure Chemical, Osaka, Japan) at a final concentration of 100 mM, followed by incubation at room temperature for 3 days. Primary antibodies against E-cadherin (rabbit polyclonal, Cat#20874-1-AP), N-cadherin (rabbit polyclonal, Cat#22018-1-AP), vimentin (rabbit polyclonal, Cat#10366-1-AP), maternal embryonic leucine zipper kinase (MELK; recombinant rabbit monoclonal, Cat# 82893-2-RR), and glyceraldehyde-3-phosphate dehydrogenase (GAPDH; rabbit polyclonal, Cat# 10494-1-AP) were purchased from Proteintech (Rosemont, IL, USA). An HRP-conjugated anti-rabbit IgG secondary antibody was purchased from Promega (Cat# W4011). Recombinant TGF-β1 (produced by HEK293 cells) was purchased from PeproTech (Cranbury, NJ, USA). OTSSP167, the MELK inhibitor, was purchased from MedChemExpress (Monmouth Junction, NJ, USA).

The human lung adenocarcinoma cell line A549 (RCB0098) was provided by Riken BRC (Ibaraki, Japan) through the National BioResource Project of the MEXT/AMED, Japan. The cells were maintained in Dulbecco’s modified Eagle’s medium (D-MEM) supplemented with 10% (v/v) fetal bovine serum (FBS; Merck, Darmstadt, Germany) and an antimicrobial cocktail (100 units/mL penicillin G, 100 µg/mL streptomycin; Fujifilm Wako Pure Chemical). Subconfluent cells were detached using 0.05% (w/v) trypsin/0.53 mM EDTA solution (Fujifilm Wako Pure Chemical), followed by centrifugation (300 × *g*, 3 min). The pellet was resuspended in fresh culture medium and used for subsequent experiments.

### 2.2. Inductively coupled plasma-optical emission spectrometry (ICP-OES)

Atelocollagen acid solution was mixed with TP(DMSO) at final concentrations ranging from 50 µM to 800 µM in 5 mM HCl aq. and incubated at 4°C for 3 days to allow cross-linking. The reaction mixtures were subjected to ultrafiltration using Amicon® Ultra Centrifugal Filter with a 3 kDa molecular weight cut-off filter (Merck). The resulting filtrates were diluted 1:50 with deionized water and analyzed using an ICP atomic emission spectrometer ICPE-9820 (Shimadzu, Kyoto, Japan). Platinum concentrations were determined from a standard curve generated using a platinum standard solution (Fujifilm Wako Pure Chemical).

### 2.3. Determination of cross-linking sites by liquid chromatography–mass spectrometry (LC-MS)

Bovine collagen (at a final concentration of 1 mg/mL) was mixed with TP(DMSO) at final concentrations ranging from 10 µM to 800 µM in 0.1 M acetic acid, and the cross-linking reaction was carried out at 4°C for 16 h. Following incubation, the reaction mixture was extensively dialyzed against 0.1 M acetic acid at 4°C using a Slide-A-Lyzer MINI dialysis unit (10 K MWCO; Thermo Fisher Scientific, Waltham, MA, USA). To determine the amino acid composition and the collagen concentration, which was calculated by summing the amino acid contents, the samples were subjected to amino acid analysis using an L-8900 amino acid analyzer (Hitachi, Tokyo, Japan) after gas-phase acid hydrolysis (6 M HCl/1% (v/v) phenol at 110°C for 20 h under N_2_) [22].

For the determination of cross-linking sites, the TP(DMSO)-reacted samples (8 µg) were denatured at 60°C for 30 min in a digestion buffer (100 mM Tris-HCl, 1 mM CaCl_2_, pH 7.6) and then digested with trypsin (enzyme/substrate ratio of 1:50) at 37°C for 16 h. The resulting digests were subjected to LC-quadrupole time-of-flight (QTOF)-MS analysis. The MS scan and MS/MS acquisition were performed using an ultra-high resolution QTOF mass spectrometer (maXis II, Bruker Daltonics, Bremen, Germany) coupled to a Prominence UFLC-XR system (Shimadzu). Peptides were separated using an Ascentis Express C18 column (2.7 µm particle size, 150 mm × 2.1 mm; Merck), as described previously [23]. Acquired MS/MS spectra were searched against the UniProtKB/Swiss-Prot database (release 2024-05-29). Peptide fragments identified with ≥95% confidence in the control sample (without TP(DMSO) treatment) were chosen for quantification. For sequences yielding multiple fragments due to differences in post-translational modifications or enzymatic digestion, representative fragments were selected for analysis based on their detection sensitivity. The relative abundance of each peptide across TP(DMSO) concentrations was calculated from the peak area ratios of monoisotopic extracted ion chromatograms (mass precision range = ± 0.01) of the precursor MS ions.

### 2.4. Mechanical testing

For compressive testing, 800 µL of gels containing atelocollagen at concentrations of 2 mg/mL, 3 mg/mL, or 4 mg/mL, TP(DMSO) at 0, 50, 100, 200, or 400 µM, 30 mM phosphate buffer (pH 7.4) and 120 mM NaCl were prepared in a 24-well plate (Watson, Tokyo, Japan). The gel precursor solutions were dispensed into the wells and incubated at 37°C for 3 h. Force-displacement curves were obtained using a universal material testing machine UCT-500 (A&D, Tokyo, Japan) equipped with a 10 N load cell and a cylindrical probe (8 mm diameter) under constant velocity pushing at 10 mm/min. The force-displacement curves were converted to force-strain curves based on the gel thickness (approximately 4 mm). Compressive elastic modulus was calculated as the slope of the elastic linear region.

For rheological measurements, 2 mL of gels containing 2 mg/mL atelocollagen, TP(DMSO) at 0, 50, 100, 200, or 400 µM, 30 mM phosphate buffer and 120 mM NaCl were prepared by casting the precursor solutions into silicone molds (35 mm diameter, 2 mm height) and incubating at 37°C for 3 h. The resulting gel was transferred to the sample stage of a rheometer MCR 302 (Anton Paar, Graz, Austria). Storage elastic modulus (G′), loss elastic modulus (G″), and complex modulus (G*) were recorded under constant oscillatory shear at 1 Hz with increasing strain (0.01–100%) at 25°C.

### 2.5. Preparation of gel substrates and on-gel cell culture

For preparation of atelocollagen coated substrates, 250 µL of atelocollagen solution (50 µg/mL) in 30 mM phosphate buffer containing 120 mM NaCl was dispensed into each well of a 24-well plate and incubated at 37°C for 1 h. The substrates were rinsed with the culture medium prior to cell seeding. A549 cells were resuspended in D-MEM containing 1% (v/v) FBS and seeded onto the substrates.

For time-lapse imaging, cells were resuspended with D-MEM containing 1% (v/v) FBS and seeded onto the substrates. The culture plate was placed in a confocal microscope BC43 (Oxford Instruments, Oxford, UK) equipped with a CO_2_ supply and temperature control system. Time-lapse images were acquired for 18 h. Cell migration trajectories and migration distances were analyzed for 15 randomly selected cells per field of view using ImageJ software.

### 2.6. Cell proliferation assay

Substrates were prepared as described above. Cells were resuspended in D-MEM containing 10% (v/v) FBS and seeded onto the substrates. After the designated culture period, the medium was replaced with fresh medium containing Hoechst 33342 (Dojindo Laboratories, Kumamoto, Japan) to stain cell nuclei. Five fluorescence images per well were acquired using a fluorescence microscope BZ-X1000 (Keyence, Osaka, Japan). The number of stained nuclei was quantified and converted to cell density (cells/mm^2^) using ImageJ software. Quantitative analysis of cell aggregation was conducted using the same images at day 3. The positions of stained nuclei were measured (>200 per view), and aggregated cells were determined as cells with at least one neighboring nucleus within 10 µm.

### 2.7. Western blotting

A549 cells were cultured on the substrates in D-MEM containing 1% (v/v) FBS for 2 days, as described above. For plastic dishes, including a collagen-coated dish, the culture medium was removed, and cells were directly lysed with radioimmunoprecipitation assay (RIPA) buffer (Nacalai Tesque, Kyoto, Japan). For gel matrix samples, the medium was removed, and gels were treated with 0.01% (w/v) recombinant collagenase (NP-collagenase; Nippi, Tokyo, Japan) in phosphate-buffered saline (PBS; Toho, Tokyo, Japan) and incubated for 10 min to detach the cells. The detached cells were collected into a 1.5 mL-tube and centrifuged at 800 × *g* for 3 min. The resulting cell pellets were lysed with RIPA buffer.

Protein concentrations were determined using a bicinchoninic acid (BCA) protein assay kit (Nacalai Tesque). Sodium dodecyl sulfate (SDS) samples were prepared using 4× SDS sample buffer (Fujifilm Wako Pure Chemical, Osaka, Japan). The samples were separated by 9% (w/v) polyacrylamide gel electrophoresis and transferred onto polyvinylidene difluoride (PVDF) membranes according to the standard procedures.

Membranes were blocked with 1% (w/v) bovine serum albumin (BSA; Fujifilm Wako Pure Chemical) in PBS for 1 hour and then incubated overnight with primary antibodies diluted in 0.1% (w/v) BSA/PBS (1:1000 for E-cadherin, N-cadherin, vimentin, and MELK; 1:5000 for GAPDH). After washing three times with PBS containing 0.05% (v/v) Tween 20 (PBST), membranes were incubated with HRP-conjugated secondary antibody (1:5000) for 30 min. Following three washes with PBST, immunoreactive bands were visualized using a chemiluminescence detection system (Chemi-Lumi One series; Nacalai Tesque) and imaged with a G:BOX Chemi XRQ system (Syngene, Bengaluru, India).

All experiments were performed in triplicate, and band intensities were quantified using ImageJ software.

### 2.8. Gene expression analysis

Cells were cultured on the indicated substrates for 2 days, as described above. For cells cultured on gel matrices, cells were harvested by incubation with a collagenase solution for 10 min to detach the cells, followed by transfer of the cell suspension to 1.5 mL tubes and centrifugation at 800 × *g* for 3 min. For cells cultured on collagen-coated substrates, cells were directly subjected to the following treatment. The resulting cell pellets were subjected to total RNA isolation using the NucleoSpin RNA Plus kit (Takara Bio, Shiga, Japan) according to the manufacturer’s instructions.

DNA microarray analysis was performed by Filgen (Aichi, Japan) using the GeneChip™ Clariom S Assay (Thermo Fisher Scientific). Pathway enrichment analysis was conducted using the R programming language with reference to the KEGG pathway database.

For quantitative reverse transcription PCR (qRT-PCR), total RNA solutions were prepared as described above. Gene-specific primers for *MELK* (forward: 5′-CCAAGAAGGCTCGGGGAAAA-3′; reverse: 5′-GAATGGGGTAGCACTGGCTT-3′) and *GAPDH* (forward: 5′-AGCCGCATCTTCTTTTGCGT-3′; reverse: 5′-GCCCAATACGACCAAATCCGT-3′) were synthesized by Eurofins Genomics (Tokyo, Japan). Total RNA, primer solutions, and Thunderbird™ Next SYBR™ qPCR Mix (Toyobo, Osaka, Japan) were mixed according to the manufacturer’s instructions, and amplification curves were recorded using a QuantStudio™ 3 real-time PCR system (Thermo Fisher Scientific). Relative mRNA expression levels were calculated using the ΔΔCt method, with GAPDH used as an internal control.

### 2.9. Anticancer drug sensitivity assay

A549 cells were cultured on collagen-coated substrates, fibrillar collagen gels, or platinum cross-linked collagen gels containing 400 µM TP(DMSO) for 2 days to induce mechanoresponsive genomic alterations. The culture medium was then replaced with fresh D-MEM containing 1% (v/v) FBS in the absence or presence of OTSSP167 at concentrations ranging from 10 to 1000 nM. After an additional 1 day of culture, live, dead, and total cells were differentially stained by adding calcein-AM, propidium iodide (PI), and Hoechst 33342 to the culture medium, followed by incubation for 30 min.

Fluorescence images were acquired using BZ-X1000. Cell viability was evaluated by calculating the ratio of PI-stained area to Hoechst 33342–stained area. Stained areas were quantified using ImageJ software.

### 2.10. Statistical analysis

Statistical analyses were performed for data of cell migration distance (Figure 3E), quantitative western blot (Figure 4), and cytotoxicity (Figure 6C) using the statistical software JMP. Statistical significance was evaluated using one-way analysis of variance (ANOVA), followed by appropriate post hoc tests for multiple comparisons. A significance level of p < 0.05 was used to determine statistical significance, and p values for statistically significant differences are indicated in the corresponding figures.

## 3. Results and discussion

### 3.1. Determination of the number and sites of platinum cross-linking sites on collagen

In this study, TP(DMSO) complex was used as a cross-linker for collagen. To determine whether the degree of collagen cross-linking could be controlled by varying the amount of added platinum, collagen-unbound platinum was quantified by ICP-OES after the cross-linking reaction (Figure 1A). The amount of collagen-bound platinum increased linearly between 50 µM and 400 µM TP(DMSO) and reached a plateau at 400 µM and above. The number of bound platinum complexes per collagen molecule (trimer) was calculated from the ratio of platinum to collagen concentration, yielding a maximum number of approximately 13 (Figure 1B).

**Figure 1.**
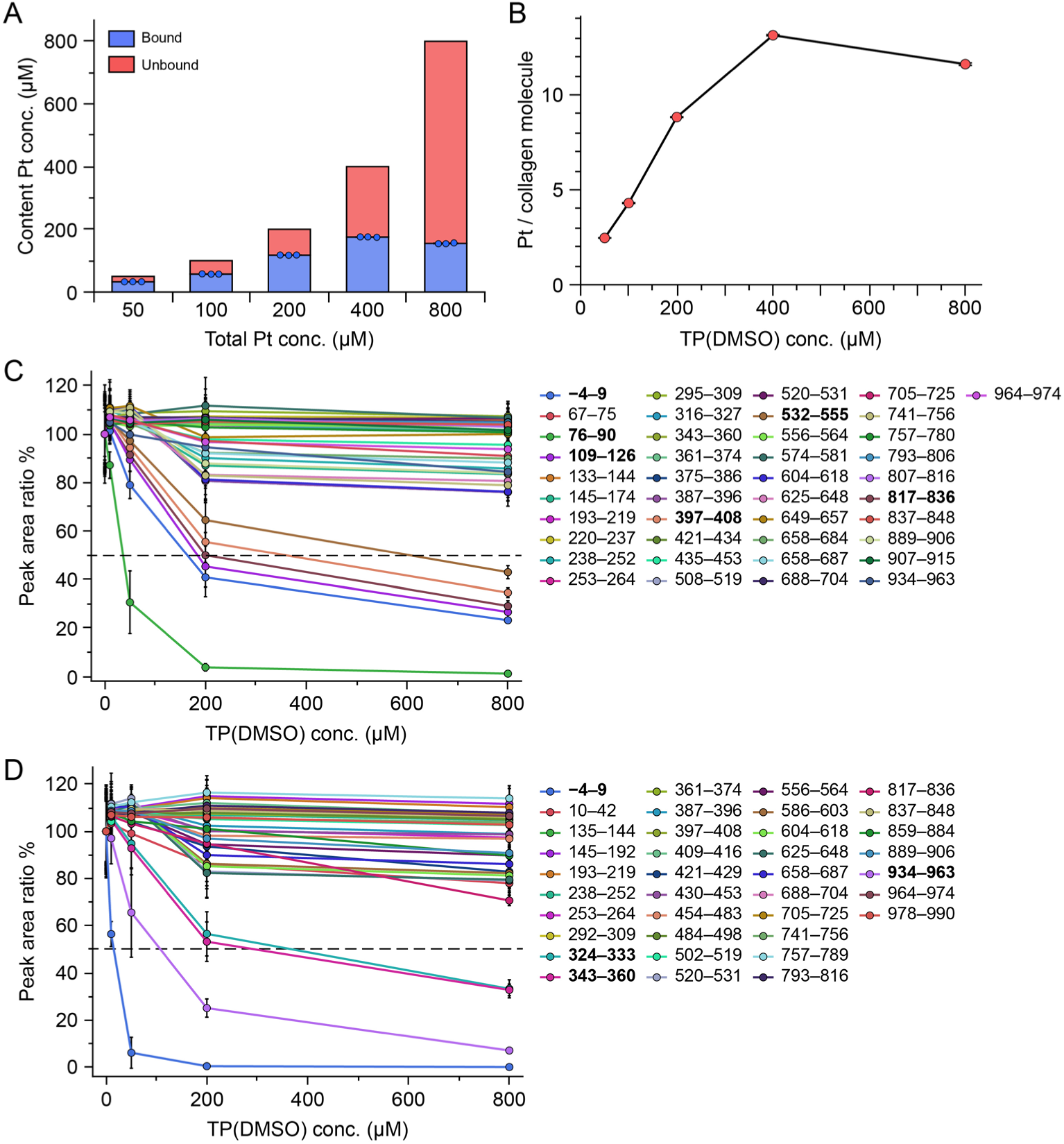
Quantification of collagen-bound platinum complexes and analysis of their cross-linking sites. (A) Platinum content in collagen-bound and -unbound fractions. Atelocollagen solutions (4 mg/mL) were incubated with TP(DMSO) at varying concentrations indicated, followed by ultrafiltration. The amount of unbound platinum in the filtrate was quantified by ICP-OES. Collagen-bound platinum concentration was calculated as the difference between total and unbound platinum concentration. (B) Estimated number of platinum bound per trimeric collagen molecule. The number was calculated from the concentration of collagen (4 mg/mL, molecular weight: 300 kDa) and collagen-bound platinum. Data represent mean ± standard error (n = 3). (C, D) Identification of platinum cross-linking sites by LC-MS analysis. Bovine type I collagen (1 mg/mL) was cross-linked with varying concentrations of TP(DMSO) in 0.1 M acetic acid, followed by trypsin digestion. Peak intensities of tryptic peptides from α1 (C) and α2 (D) chains detected with ≥95% confidence were compared across the TP(DMSO) concentrations. Sequences with a decrease of greater than 50% in peak area are shown in bold. All analyzed sequences are listed in Table S1 and S2. Data represent mean ± standard deviation (n = 3).

Platinum-mediated cross-linking sites in type I collagen were further investigated by comprehensive detection of tryptic peptides using LC-MS (Figure 1C and D). As a result, six peptides in the α1 chain and four peptides in the α2 chain showed decreases upon platinum cross-linking. Considering that a collagen molecule is composed of two α1 chains and one α2 chain, leading to at least 16 potential cross-linking sites within a collagen molecule. To identify amino acid residues which serve as ligands for TP(DMSO), amino acid analysis was performed following cross-linking (Figure S1). The abundance of sulfur-containing amino acids, including Met and Cys, decreased after cross-linking, indicating that TP(DMSO) binds to the side chains of these amino acid residues. Consistently, all peptides identified by LC-MS analysis contained at least one Met residue, with the exception of one peptide 934–963 in α2 chain. Notably, the peptide 934–963 includes His-rich region, suggesting His residue is an additional cross-linking site.

The triple-helical region of porcine collagen contains 19 Met and 16 His residues, both of which are potential coordination sites for TP(DMSO) [21]. In our previous study, the number of collagen-binding platinum complexes was found to be similar to the value shown in Figure 1B, even when the concentration of collagen or atelocollagen was different. This result suggests that TP(DMSO) binds to collagen in a stoichiometric manner. Assuming that TP(DMSO) functions as a divalent cross-linker, the theoretical number of bound platinum complex per collagen molecule is 17.5, which is reasonable for the quantitative results. The LC-MS analysis supports this estimation (Figure 1C and D). All peptides containing Met or His residues, with the exception of His422 in the α2 chain, were identified as potential platinum cross-linking sites. In amino acid analysis following cross-linking, His residues were detected predominantly in their intact form (Figure S1). Nevertheless, this may be attributable to dissociation of platinum-bound histidine due to ligand exchange with chloride ions during heating in concentrated hydrochloric acid. Taken together, these results strongly suggest that Met and His residues within the triple-helical region serve as the primary cross-linking sites for TP(DMSO) in collagen, whereas Cys residue in telopeptides may also act as minor cross-linking sites in collagen retaining telopeptides.

Interestingly, peptides at positions 76–90 in the α1 chain and 4–9 in the α2 chain efficiently and similarly decreased in the LC-MS analysis according to cross-linking, suggesting platinum cross-linking between the regions. The 76–90 sequence includes a multifunctional sequence MKGHRGF, which interacts with heparan sulfate [24,25], pigment epithelium-derived factor (PEDF) [26], and telopeptides, which are non-triple-helical regions at the termini of triple-helical domain and important for fibril formation of collagen [27,28]. Notably, the MKGHRGF sequence has been reported to be typically cross-linked *in vivo* with the C-terminal telopeptide of the α1 chain [28]. Therefore, TP(DMSO) was likely to effectively cross-link methionine residues within MKGHRGF and N-terminal telopeptide of the α2 chain, thereby its inhibitory effect on collagen fibril formation in the normal fashion.

### 3.2. Characterization of tunable mechanical properties of PCG

Based on the dynamic range of TP(DMSO) concentration for collagen-binding, the controllability of mechanical properties was assessed by measuring the elastic moduli of PCGs. Compressive elastic modulus was first evaluated by conventional compressive testing (Figure 2A). The maximum compressive modulus was observed at TP(DMSO) concentration of 200 µM, 200 µM, and 100 µM for atelocollagen concentration of 4 mg/mL, 3 mg/mL, and 2 mg/mL, respectively. This result indicated that excessive TP(DMSO) did not enhance the compressive modulus and, rather, tended to suppress its increase. However, the perceived stiffness of PCGs containing 2 mg/mL atelocollagen did not qualitatively correspond to the compressive modulus values (Supplemental video 1). Therefore, shear elastic modulus of the gels was additionally evaluated by rheometric analysis to capture another aspect of the mechanical properties (Figure 2B and S2). In contrast to the compressive modulus, the shear elastic modulus increased according to TP(DMSO) concentration.

**Figure 2.**
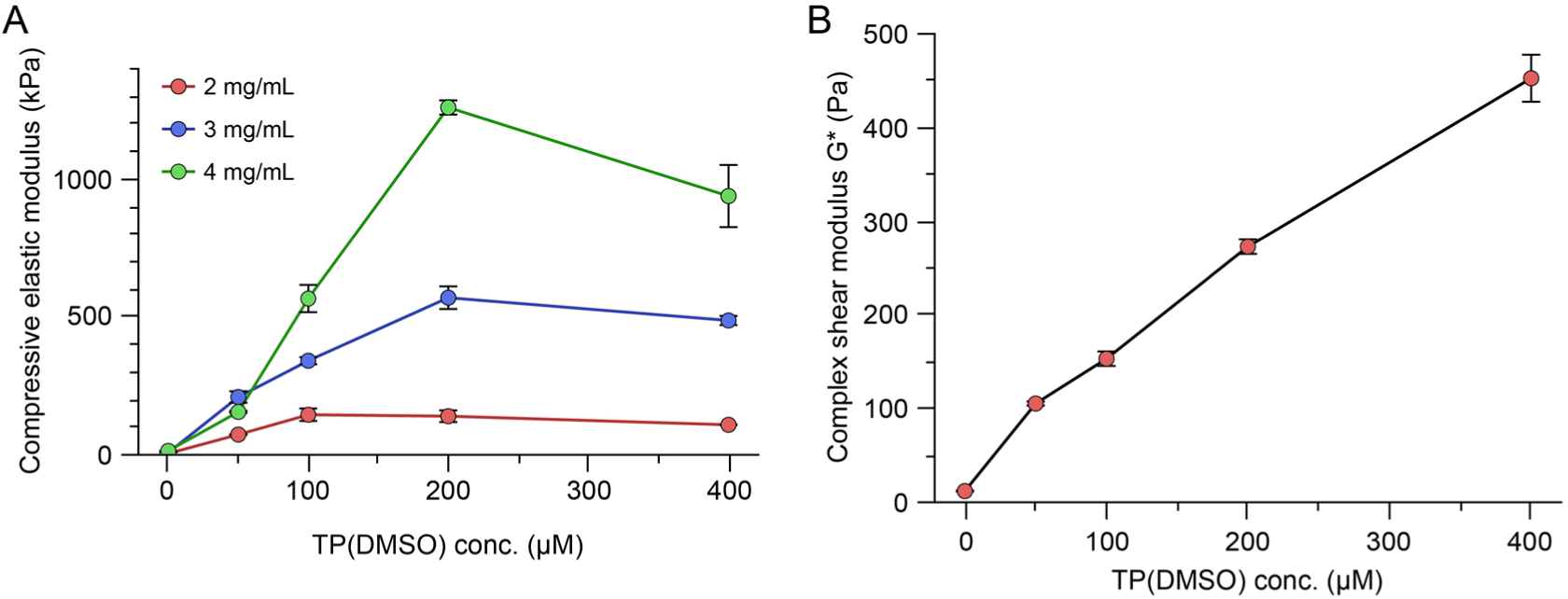
Mechanical characterization of PCGs by compressive and rheological measurements. (A) Compressive elastic modulus of gels with varying concentrations of atelocollagen and platinum complex. Gels were prepared by mixing atelocollagen with TP(DMSO) at the indicated concentrations. (B) Complex shear elastic modulus of gels with varying concentrations of platinum complex. Gels were prepared by mixing varying concentrations of TP(DMSO) and atelocollagen at fixed final concentration of 2 mg/mL. The complex shear modulus was measured using a rheometer. Data represent mean ± standard error (n = 3).

The compressive modulus of the gels did not show a similar pattern to the amount of collagen-binding platinum (Figure 1B and 2A), suggesting that excess platinum may partially mask potential cross-linking sites on collagen molecules, thereby limiting network formation and suppressing the increase in compressive modulus. In contrast, the shear modulus exhibited a trend that closely paralleled the pattern shown in Figure 1B. This finding was also consistent with the qualitative perception of the gels (Supplemental video 1). During PCG formation, both platinum cross-linking and fibril formation, which is intrinsic to native collagen fibrillogenesis, contribute to the resulting physical strength [20]. Hence, the shear modulus likely exhibited the combined effects of these processes.

### 3.3. A549 cell responses to PCGs with tunable mechanical properties

Here, the applicability of PCG with tunable mechanical properties to cell biological studies was evaluated using a mechanoresponsive EMT model of A549 cells. Collagen concentration was fixed at 2 mg/mL because the PCG series almost covered the physiological range of compressive modulus in most soft tissues within the dynamic range [29,30]. Prior to the subsequent assessments, the effects of free platinum complexes eluted from PCGs on cellular functions were evaluated (Figure S3). No significant alterations were observed in cell viability, and only minor variations in gene expression were detected, which were not considered to affect biologically meaningful gene sets. Morphological differences were examined for cells cultured on a plastic dish (PS), a conventional atelocollagen fibrillar gel (FG), or PCGs with varying mechanical properties (Figure 3A). On PS, cells displayed a cobblestone-like epithelial morphology, whereas treatment with TGF-β1 induced EMT, resulting in scattering and fibroblast-like morphology. Cells cultured on an atelocollagen-coated dish (Col coat) exhibited a spiky shape, which was different from that on PS regardless of TGF-β1 treatment. On FG, which has the lowest mechanical parameters among the gel matrices, cells formed isolated colony-like clusters. PCGs induced morphological changes that depended on TP(DMSO) concentration. On PCG containing 50 µM TP(DMSO) (PCG(50)), cells tended to aggregate; however, increasing the amount of TP(DMSO) promoted scattering and yielded morphologies resembling those observed on Col coat (Figure 3B and S4). These phenotypes were clearly different from those on FG.

**Figure 3.**
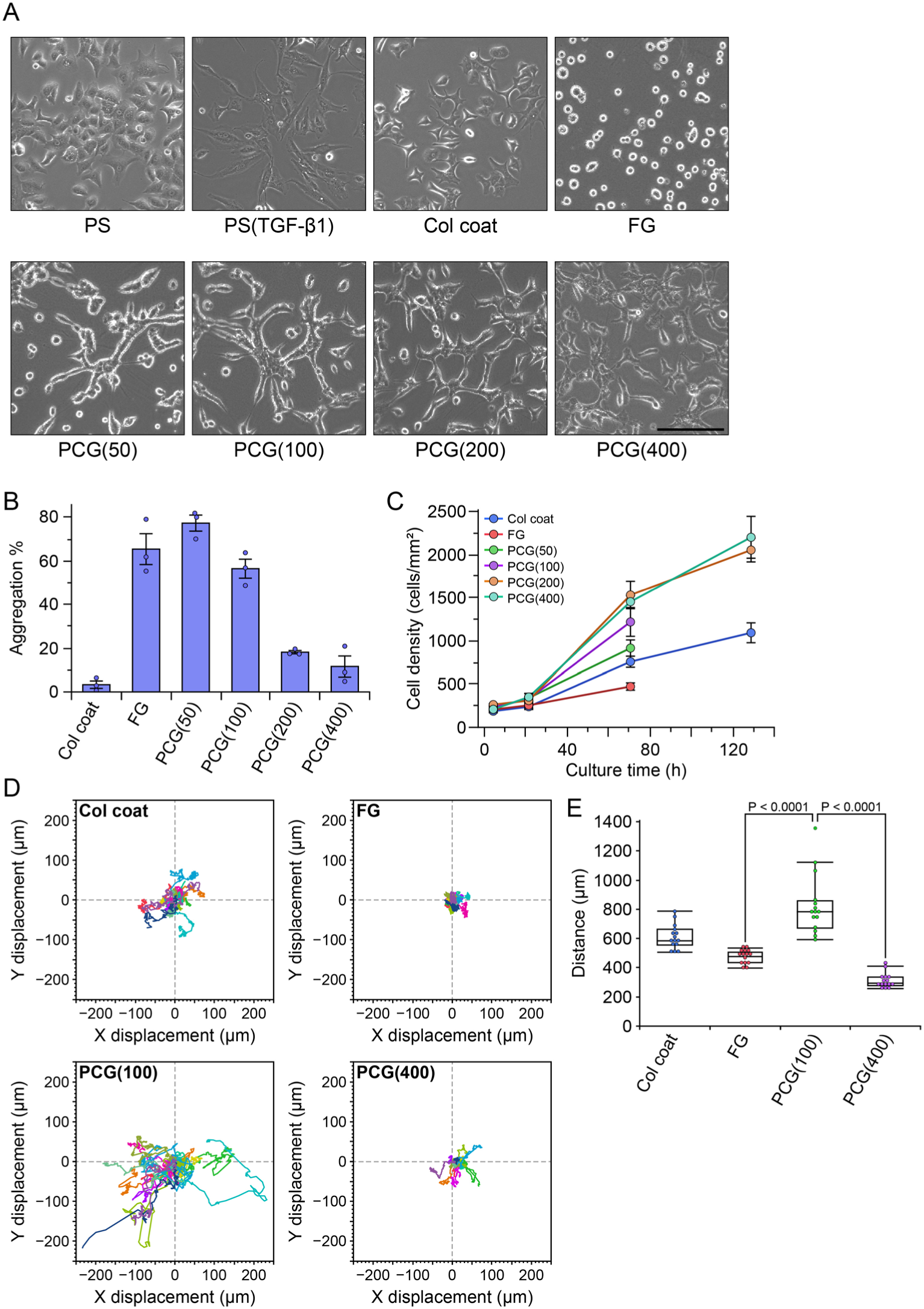
A549 cell responses to different substrates. (A) Cell morphology on various substrates. A549 cells were cultured on plastic dishes (untreated, TGF-β1-treated [PS(TGF-β1)], or atelocollagen-coated [Col coat]), conventional atelocollagen fibrillar gel (FG), or PCGs containing TP(DMSO) at the indicated concentrations (µM). All gels are composed of 2 mg/mL collagen. Images were acquired after 2 days of culture. Scale bar represents 200 µm. (B) Quantification of cell aggregation. Cells were cultured on the substrates for 3 days, stained with Hoechst 33342, and imaged using a fluorescence microscope. Aggregated cells were defined as cells with at least one neighboring nucleus within 10 µm. Bars and error bars represent the mean ± standard error (n = 3). (C) Cell proliferation on different substrates. Cells were cultured for the indicated periods and stained with Hoechst 33342. Five images were acquired per well. The number of nuclei was quantified and converted to cell density (cells/mm^2^). Quantification for FG, PCG(50), and PCG(100) was terminated at day 3 because individual cells could not be distinguished due to extensive aggregation. Dots and error bars represent the mean ± standard deviation. (D) Cell migration trajectories. Trajectories of 15 cells randomly selected per condition were tracked from time-lapse images recorded for 18 h. (E) Quantification of migration distance. Total migration distance for each cell was measured from the trajectories shown in panel D. Statistical analysis was performed using ANOVA followed by the Steel-Dwass post hoc test.

The effects of gel mechanical properties on cell proliferation were subsequently examined (Figure 3C). Among the substrates, FG exhibited the lowest proliferation activity, followed by Col coat, and then PCGs. Within the PCG series, cell proliferation activity increased with increasing TP(DMSO) concentration and reached comparable levels on PCG(200) and PCG(400). Notably, although cells cultured on Col coat were almost confluent at day 5, similar to those on PCG(200) and PCG(400), cells on PCGs proliferated more densely, resulting in approximately two-fold higher cell numbers than on Col coat (Figure S5). Interestingly, cells on PCG(50) and PCG(100) further aggregated to form large multicellular clusters at day 5.

Differences in the motility of A549 cells on substrates with varying mechanical properties were additionally evaluated (Figure 3D, E). Cells exhibited distinct migration behaviors on Col coat, FG, PCG(100), and PCG(400). In particular, cells on FG showed minimal displacement in one day, whereas cells on PCG(100) displayed markedly enhanced motility compared with those on other substrates. These results indicate that active migration of A549 cells requires an intermediate substrate stiffness. Furthermore, pseudopodia on PCG(400) appeared to attach strongly to the substrate in the process of cell migration (Supplemental video 2), suggesting that the tight interaction between a cell and a substrate likely hinders cell motility.

Relationships between cell morphology and mechanical properties of matrices have been widely reported. A549 cells have been shown to become more isolated and spread with enhanced focal adhesion in response to increasing matrix compressive modulus [31]. Conversely, the cells tend to form aggregates and exhibit limited elongation of pseudopodia on a matrix with lower compressive modulus. Consistent with the previous observation, the cells displayed similar morphological features on the substrates with different mechanical properties (Figure 3A and B). However, although the compressive modulus of PCGs reached a plateau at TP(DMSO) concentration of 100 µM and more (Figure 2A), cell shape continued to change beyond this point. In addition, correlation coefficients between aggregation values shown in Figure 3B and the compressive and shear moduli presented in Figure 2 were −0.54 and −0.90, respectively. These results suggest that morphological cell behavior is more strongly governed by shear elasticity of the matrix.

A previous study reported that NMuMG epithelial cells exhibited morphologies similar to those observed on PCG(400) when cultured on matrices with lower loss elastic modulus [32]. In contrast, the cells formed aggregates without pseudopodia on matrices with higher loss modulus despite comparable storage modulus. Although this report provides important insight into the role of viscoelastic properties in cell behavior, the storage moduli of PCGs used in the experiments were different each other. Therefore, a more systematic investigation using PCGs with varying compressive, storage, and loss moduli would allow us to elucidate contributions of these mechanical parameters on cell behavior.

It has been reported that proliferation activity is positively regulated by substrate stiffness in different cell types [33–35]. A549 cells also exhibit enhanced proliferation on stiffer matrices [36]. In this previous study, a stiffness-tunable collagen matrix cross-linked with genipin was employed, representing a system similar to that used in the present study. As shown in Figure 3C, cell proliferation on the gel matrices showed a slightly stronger correlation with the compressive modulus than with the shear modulus shown in Figure 2 (correlation coefficients: 0.91 for compressive modulus and 0.86 for shear modulus). Although Col coat has the highest elastic modulus among all tested materials, cell proliferation on Col coat was lower than that on PCGs. This may be attributable to more extensive cell attachment compared with that on PCGs (Figure S5), which allows the substrate surface to be covered by fewer cells and may lead to contact inhibition. Conversely, PCGs may function as substrates that enable tighter cellular organization. However, cell-matrix interactions may become relatively weaker than cell-cell interactions, when the shear modulus is smaller than a certain threshold, thereby resulting in unstable adhesion to the substrate. These observations suggest distinct functions of compressive and shear elasticity of the matrix as regulators of cellular physiology.

A549 cell motility has been also investigated using collagen-coated plastic dishes or polydimethylsiloxane (PDMS)-based elastomers as culture substrates [37]. It was reported that cell motility was enhanced on substrates with lower compressive modulus. However, cell motility on FG, which was used as a substrate with the lowest stiffness, was relatively lower than other substrates. This discrepancy suggests that cells sense multiple aspects of mechanical properties of matrices; not only compressive modulus but also shear modulus and other mechanical parameters, as well as their relative balance.

Although the discussion above has focused on the mechanical properties of the matrix, it is also important to consider the effects of possible cross-linking at collagen receptor-binding sites. As shown in Figure 1C, the MKGHRGF sequence in the α1 chain (76–92) is a highly cross-linked region and also serves as a binding site for syndecan, a heparan sulfate proteoglycan. In addition, the GVMGFP sequence in the α1 chain (400–405) is a ligand for the discoidin domain receptors (DDRs). Since we have not investigated how cross-linking at the sites affects the binding of these receptors or their signal transduction, the possibility could not be excluded that cell behavior was altered by shutting down collagen receptor signaling due to cross-linking.

### 3.4. Expression of EMT-related proteins in PCGs with varying mechanical properties

To investigate whether the differences in cell morphology and motility observed on substrates with varying mechanical properties were attributed to EMT caused by the substrates, the expression levels of EMT-related proteins were analyzed (Figure 4). As a result, no significant changes were detected in the expression levels of E-cadherin and vimentin, which are upregulated upon EMT, and N-cadherin, which is downregulated during EMT, among the substrate conditions, except for PS(TGF-β1) as an EMT positive control. This result indicates that the substrate stiffness did not induce EMT in the present experimental system. Conversely, the observed differences in cell morphology and motility were therefore suggested to be regulated independently of EMT.

**Figure 4.**
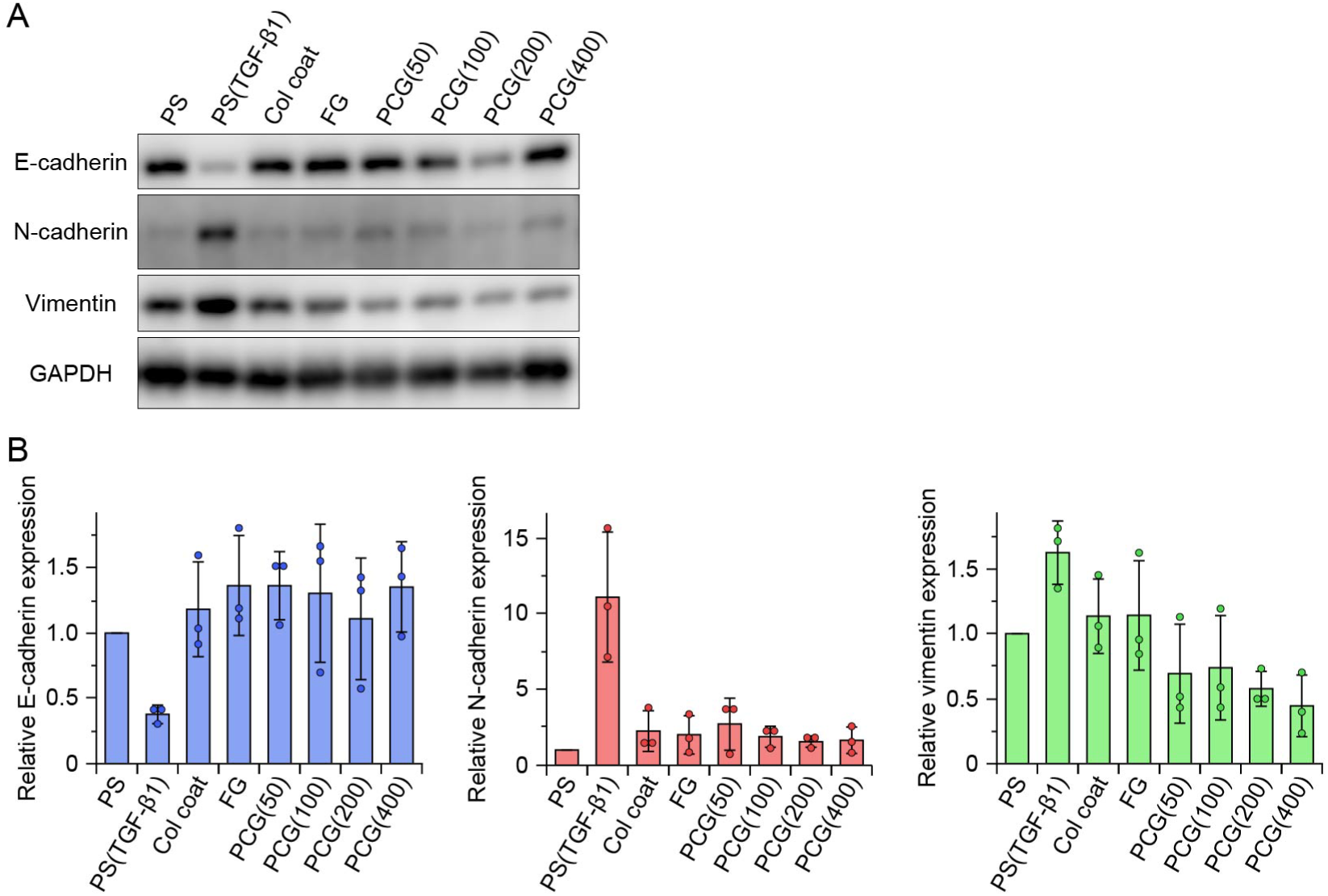
Quantitative analysis of EMT-related protein expression in A549 cells. (A) Representative western blot images of EMT-related proteins. (B) Densitometric analysis. Band intensities were quantified and normalized to the housekeeping protein GAPDH, and all values were further normalized to those of PS. Bars and error bars represent mean ± standard deviation (n = 3). Statistical analysis was performed using ANOVA followed by the Steel post hoc test.

Although various studies have examined the effects of collagen on cancer EMT [14,15], the findings have not been consistent. Even among studies restricted to A549 cells, discrepant results have been reported. For example, Shintani *et al*. demonstrated that EMT was promoted on collagen-coated dishes [18], whereas Fujisaki *et al*. were unable to reproduce this observation [19]. Interpretation is further complicated by the fact that molecular hallmarks of EMT do not necessarily occur in a coordinated manner. Fujisaki *et al*. also reported that cadherin switching occurred on a collagen gel without an increase in vimentin expression.

Our results did not exhibit an EMT-inducing effect of collagen. However, this finding does not directly contradict previous reports. The EMT-inducing effect of collagen previously reported appear to be clearly milder than that of TGF-β1. In addition, platinum cross-linked collagen could possess biological properties distinct from those of native collagen. DDR1 has been reported to mediate collagen-induced upregulation of N-cadherin in pancreatic cancer–derived BxPC3 cells [38]. Therefore, the possibility is not ignored that platinum cross-linking of the DDR-binding site (the GVMGFP sequence) interfered with DDR binding and downstream signaling, thereby attenuating EMT induction.

The relationship between matrix mechanical properties and EMT also warrants consideration. Shukla *et al*. reported that increased matrix stiffness alone does not induce spontaneous EMT [37], which is consistent with our results. In contrast, Sacco *et al*. demonstrated that matrix viscosity negatively regulates TGF-β-dependent EMT induction [32]. PCG may be applicable to provide a useful platform to dissect how compressive, shear elasticity, and viscosity of the matrix individually contribute to cancer EMT.

### 3.5. Analysis of gene expression modulated by mechanical parameters

Based on the above results, the differences in cell morphology and motility observed on different matrices were suggested not to be attributable to EMT. Here, to further investigate the underlying mechanisms, comprehensive gene expression analysis was performed using DNA microarrays. Considering that the platinum complexes themselves have minimal effects on gene expression (Fig. S3B), enrichment analysis revealed that many of genes associated with cell proliferation, division, and cell cycle progression were upregulated in cells cultured on PCG(400) compared with those on FG (Fig. 5A). This finding is consistent with the enhanced cell proliferation observed on PCG(400) (Figure 3C). In contrast, little change was detected in the expression of genes encoding adherens junction proteins or transcription factors involved in the mechanism of EMT. In addition, no significant downregulated pathway was found in the analysis.

**Figure 5.**
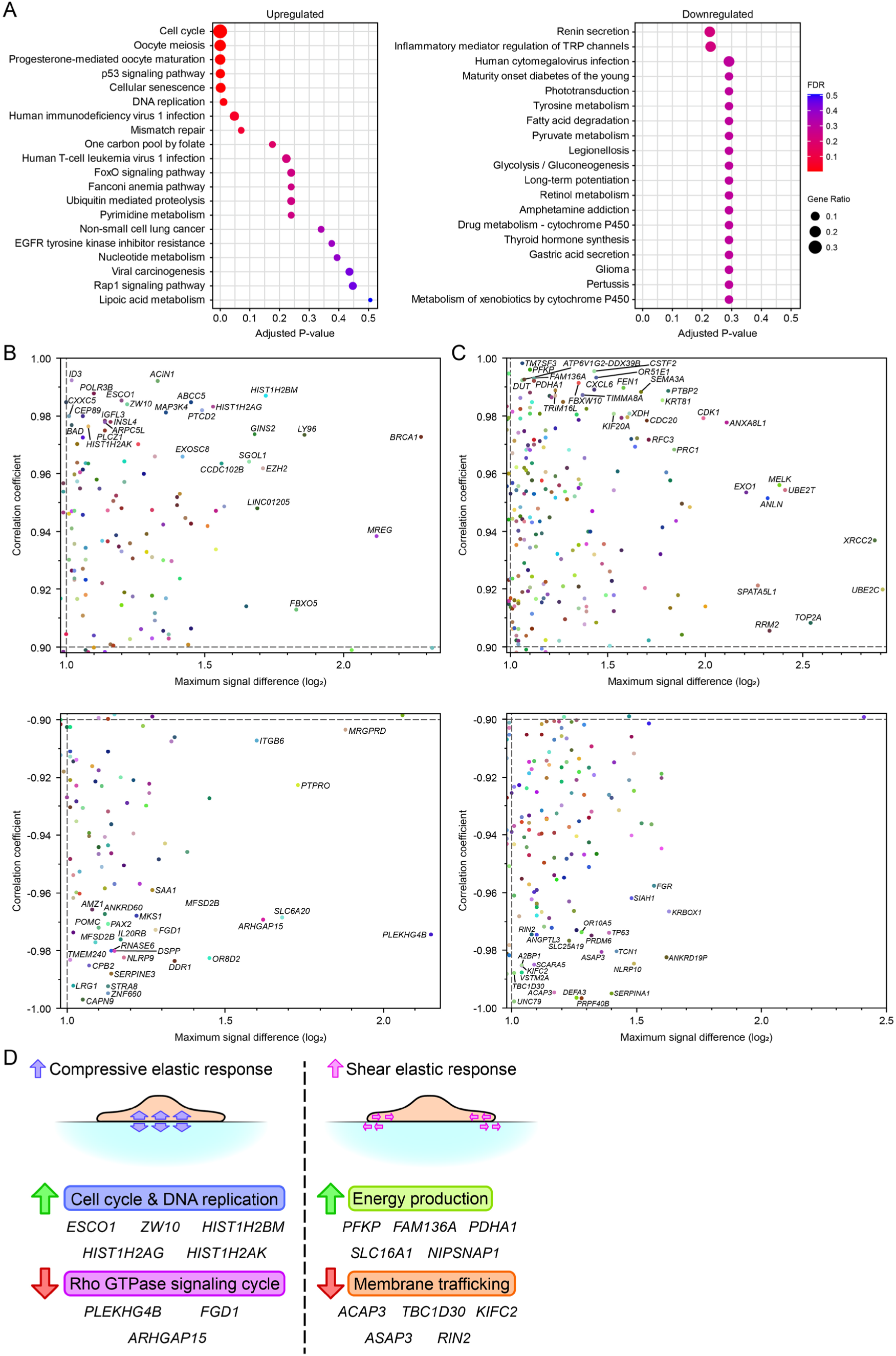
Gene expression profiling and correlation analysis of matrix mechanical properties. (A) Pathway enrichment analysis. DNA microarray data from FG and PCG(400) samples were subjected to pathway enrichment analysis using the KEGG database. Pathways upregulated or downregulated in PCG(400) relative to FG are shown. (B, C) Correlation analysis with matrix mechanical parameters. DNA microarray signal intensities obtained from FG and PCG samples were analyzed for correlations with compressive elastic modulus (B) or shear elastic modulus (C), as shown in Figure 2. Genes exhibiting large expression changes and positive or negative correlations are plotted in the upper (positive correlation) and lower (negative correlation) panels, respectively. (D) Schematic summary of correlation analysis. Representative genes associated with the indicated biological processes are shown.

**Figure 6.**
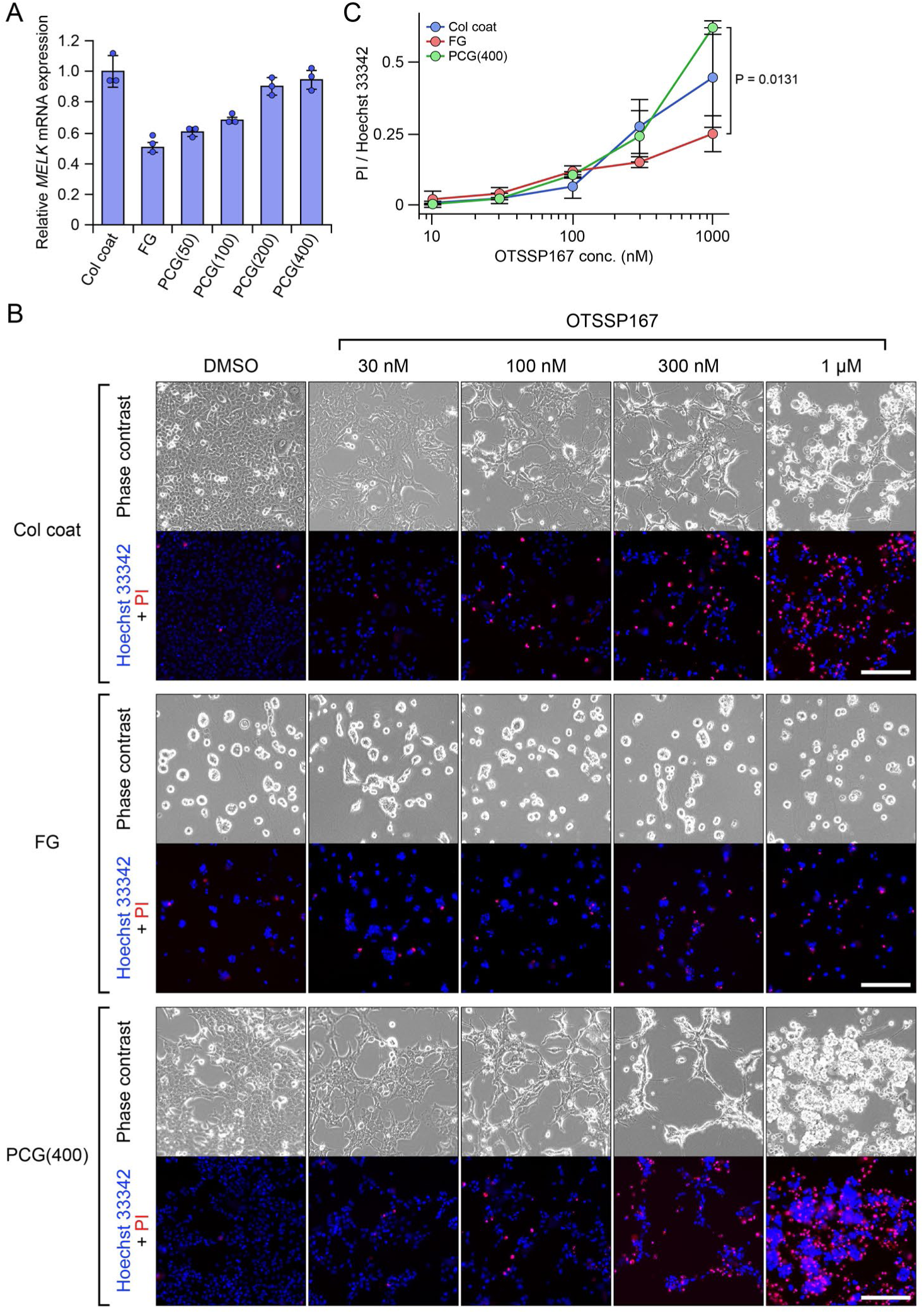
Stiffness-dependent upregulation of *MELK* expression and altered sensitivity to a MELK inhibitor in A549 cells. (A) Quantitative PCR analysis of *MELK* mRNA expression. Cells were cultured on each substrate for 2 days prior to RNA extraction. Bars and error bars represent mean ± standard deviation (n = 3). (B) Fluorescence images of A549 cells treated with the MELK inhibitor OTSSP167. After culturing on each substrate for 2 days, cells were incubated for one additional day in the presence of OTSSP167. Cell nuclei and dead cells were stained with Hoechst 33342 and PI, respectively. Scale bar represents 200 µm. (C) Quantitative evaluation of cytotoxicity. The ratio of PI-stained area to Hoechst 33342–stained area was calculated from the images shown in panel B. Dots and error bars represent mean ± standard deviation (n = 3). Statistical analysis was performed using ANOVA followed by the Tukey-Kramer post hoc test.

To identify factors whose expression was modulated by matrix mechanical properties, correlation analysis was performed between gene expression levels and the compressive or shear elastic moduli of FG and PCGs (Figures 5B and C). As a result, regardless of whether the correlations were positive or negative, the number of genes showing higher correlations with shear modulus was greater than that for compressive modulus. This indicates that gene expression was more strongly associated with shear elasticity of the matrix in the present system.

Overall, the differentially expressed genes tended to be associated with cell proliferation and cancer malignancy; nevertheless, they showed distinct trends in classified groups divided by the correlation analysis (Figure 5D). The top 20 genes with the highest absolute correlation coefficients for each elastic modulus are listed in Table S3 and S4. For compressive modulus, genes involved in cell cycle progression tended to show positive correlations, whereas genes associated with the Rho GTPases activation cycle exhibited negative correlations. In contrast, for shear modulus, genes related to metabolic pathways, such as glycolysis, showed positive correlations, while genes involved in the activation of membrane trafficking–related GTPases, including Arf and Raf, tended to show negative correlations. In addition, correlation analysis was also performed with the cell aggregation score shown in Figure 3B; yielding results approximately opposite to those obtained for shear modulus (Figure S5).

Taken together, the gene expression analysis revealed that little change was unexpectedly observed in the expression of genes associated with EMT or mechanotransduction. In contrast, gene groups involved in cell proliferation and energy production were upregulated in response to matrix mechanical parameters. Considering the relationship with cell proliferation shown in Figure 3C, essential processes for cancer progression, such as cell cycle and energy production, may be controlled by distinct mechanical drivers, namely compressive and shear elasticity.

Members of the Rho GTPase family, such as RhoA, Cdc42, and Rac1 are known to intricately control pseudopodium formation and cell motility [39,40]. As shown in Figures 3D and 3E, cell motility was highest on PCG(100), which exhibited the highest compressive modulus among the tested gels. This observation and correlation analysis, in which gene expression of RhoGAPs and RhoGEFs may be downregulated according to compressive elasticity, suggest that Rho GTPase activity is attenuated on a matrix with higher compressive modulus, thereby potentially losing cell–matrix interactions and promoting dynamic reorganization of pseudopodia. Consequently, cell–cell interactions became relatively dominant, potentially contributing to the formation of characteristic cell aggregates, as observed in Figures 3A and S4. However, the expression levels of both genes encoding FGD1 RhoGEF and ARHGAP15 RhoGAP exhibited negative correlations between compressive modulus and expression level (Table S3), have been reported to promote cancer cell migration [41,42]. Therefore, our results do not straightforwardly support the previous findings. Overall, the relationships among matrix stiffness, gene expression, and intrinsic molecular function may be regulated in a complex manner with involving multiple molecules. Therefore, further investigation will be required to clarify the mechanisms.

Activity of other small GTPase families, such as Rab and Arf, was suggested to be negatively correlated with shear elasticity of the matrix (Figure 5C and Table S4). Rab and Arf proteins are involved not only in membrane trafficking but also in actin cytoskeleton rearrangement [43,44]. Therefore, their downregulation may contribute to the formation of highly adhesive pseudopodia and the consequent suppression of cell migration on the stiffer matrix (Figure 3D, E, and Supplemental video 2). Nevertheless, ACAP3 and ASAP3, which are known as ArfGAPs, and RIN2, which is known as Rab-specific GEF, are not likely to function as consistent regulators for cell attachment and migration [45–49]. Regarding their effects on EMT, RIN2 was revealed to promote endocytosis of E-cadherin [50], which may provide a possible explanation for our observation that cadherin switching was not induced on PCGs (Figure 3). Although the biological implications of these apparent inconsistencies are currently unclear, they could reflect the complex regulation of Arf and Rab activities during cell migration or context-dependent functional changes associated with cancer progression.

DDR1 was identified as another gene whose expression was negatively correlated with compressive modulus (Figure 5B and Table S3). DDR1 is a collagen-specific receptor tyrosine kinase expressed on the cell membrane and is known to regulate cellular physiological activities in cooperation with collagen-binding integrins [51]. In addition, DDR1 has been suggested to possess mechanosensing functions [52]. Although DDR1 and its paralog DDR2 are known to be predominantly expressed in epithelial and stromal tissues, respectively, the functional differences between these receptors remain poorly understood. In epithelial cancers, it has been reported that DDR1 expression switches to DDR2 during EMT. However, the biological significance of this transition remains unclear [53].

In the present study, *DDR1* expression decreased as the compressive modulus increased, whereas *DDR2* did not exhibit a reciprocal response. This differential regulation may reflect possible distinct roles of the two DDR subtypes in cellular mechanotransduction. Nevertheless, as mentioned above, platinum complexes cross-link the GVMGFP sequence, which is the DDR-binding site. Therefore, potential alterations in DDR–collagen interactions and downstream signaling cannot be excluded. Further investigation of the interaction between platinum cross-linked collagen and DDR, as well as its impact on receptor activation and cellular behavior, may help elucidate the role of DDR signaling in cancer invasion and metastasis.

Closer inspection of the correlation analysis revealed that a variety of potential negative prognostic marker genes, not limited to lung cancer, were upregulated in association with matrix mechanical parameters. These findings suggest that a stiffer peritumoral matrix may coordinately enhance cancer malignancy. Among these genes, we focused on *MELK*, a negative prognostic marker that is highly expressed in lung adenocarcinoma and therefore conducted a more detailed analysis, as described below.

### 3.6. Influence of matrix stiffness-dependent *MELK* expression on the efficacy of a MELK-targeting antitumor drug

We first examined whether the expression of maternal embryonic leucine zipper kinase (MELK), which is known to be upregulated in lung adenocarcinoma cells, including A549 cells, and to be associated with poor prognosis [54,55], is regulated by matrix mechanical properties, as suggested by the DNA microarray analysis. *MELK* expression was quantified at both the mRNA and protein levels using quantitative PCR and western blotting, respectively (Figures 6A and S6). The qPCR results showed that *MELK* mRNA expression increased consistently with increasing shear modulus of the gel. In contrast, MELK protein expression did not exhibit a linear increase, suggesting the involvement of additional regulatory mechanisms at the level of protein synthesis and/or degradation.

We therefore hypothesized that stiffness-dependent upregulation of *MELK* expression might modulate the responsiveness of A549 cells to a MELK-targeting anticancer drug. To verify the hypothesis, cellular responses to the MELK inhibitor OTSSP167 [56] were evaluated (Figures 6B, 6C, and S7). Although the total numbers of cells were markedly different among the substrate conditions, clear differences were observed in the proportion of dead cells. The proportion of dead cells was lower on FG than on Col coat, whereas it increased on PCG(400). Quantitative evaluation of cytotoxicity was consistent with these qualitative observations.

Taken together, these results demonstrate that matrix stiffness-dependent changes in gene expression influence the responsiveness of A549 cells to the anticancer drug OTSSP167. Although MELK protein expression did not fully reflect the mRNA expression profile, sensitivity to OTSSP167 was nevertheless enhanced on stiffer matrix. This enhanced drug sensitivity is likely attributable not only to changes in MELK expression but also to alterations in related signaling pathways and associated proteins.

## 4. Conclusion

In this study, we demonstrated that a collagen cross-linked gel using a platinum complex previously developed by our group was useful as a cell culture substrate with tunable mechanical properties. In addition, the combination of a matrix series with distinct mechanical parameters, comprehensive gene expression profiling, and correlation analysis suggested that cancer cells sense the compressive and shear elasticity of the matrix and independently regulate cellular functions. This approach enabled the identification of a candidate anticancer drug. Further analysis of the cross-linking sites revealed that the platinum complex cross-linked at least one bioactive ligand for collagen receptors, such as HSPGs or DDRs, potentially resulting in the loss of collagen-specific signal transduction. Nevertheless, this system offers a versatile strategy not only for elucidating cellular mechanosensing mechanisms but also for applications in cancer diagnostics, drug discovery and therapeutic screening. Since this system can be extended to 3D as well as 2D culture platforms, we anticipate that this approach will contribute to further advances in mechanobiology research using pluripotent stem cells and patient-derived organoids.

## Acknowledgements

This work was supported by JSPS KAKENHI Grant Number JP24K15752 and by research funding provided through a collaborative research agreement with Nippi Research Institute of Biomatrix. We thank Nagaoka University of Technology Analysis and Instrumentation Center for their support and cooperation in ICP-OES measurements. Dynamic viscoelastic measurements were performed with the support of Tokyo Metropolitan Industrial Technology Research Institute (TIRI).

## CRediT authorship contribution statement

Shinichiro F. Ichise: Conceptualization, Data curation, Formal analysis, Funding acquisition, Investigation, Methodology, Project administration, Validation, Writing – original draft. Yuki Taga: Methodology, Validation. Kazumasa Fujita: Methodology, Validation. Takaki Koide: Funding acquisition, Investigation, Writing – review & editing.

## Data availability

The data that support the findings of this study are available from the corresponding author upon reasonable request.

