## Supplementary material for "Platinum Cross-linked Collagen Matrices with Tunable Stiffness as a Platform to Investigate Cellular Mechanosensing": Figure S1, Figure S2, Figure S3, Figure S4, Figure S5, Figure S6, Figure S7, Figure S8, Table S1, Table S2, Table S3, Table S4

### Supplemental information

1. Cytotoxicity assay

To prepare conditional media for assay, 1 mL of PCG(200) or PCG(400) gels were prepared in the wells of a 6-well plate, followed by addition of 4 mL of D-MEM containing 10% (v/v) FBS and incubation overnight. Collected media was added to the wells of a 96-well plate with subconfluent A549 cells. After 1 day culture, live cells were quantified using Cell Counting Kit 8 (Dojindo). Untreated culture medium was used as a control, and the cell viability was normalized using the control.

2. Quantitative PCR for investigating effect of TP(DMSO) on gene expression

A549 cells were subconfluently seeded to wells of 12-well plate. Medium was replaced with fresh culture medium or PCG(400)-treated one prepared as described above, followed by culturing overnight. Total RNA was extracted using NucleoSpin RNA Plus kit (Takara Bio). Preparation of cDNA library and DNA microarray analysis using Affymetrix Clariom S array were performed by Macrogen Japan corporation (Tokyo, Japan).

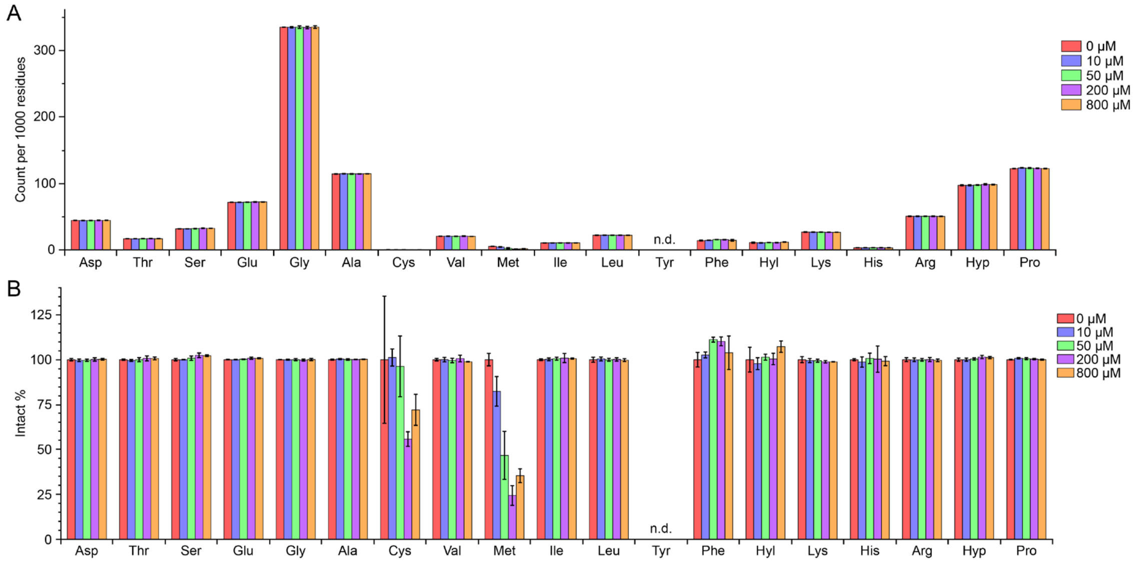


Figure S1. Characterization of amino acids binding to TP(DMSO). Bovine type I collagen (1 mg/mL) was treated with TP(DMSO) at the indicated concentrations under acidic conditions. The samples were subsequently subjected to acid hydrolysis followed by amino acid analysis. The amino acid composition is presented as (A) the number of residues per 1000 total residues and (B) values normalized to the untreated control (0 µM). Hyp indicates 4-hydroxyproline, and n.d. indicates not detected. Bars and error bars represent the mean ± standard deviation (n = 3).

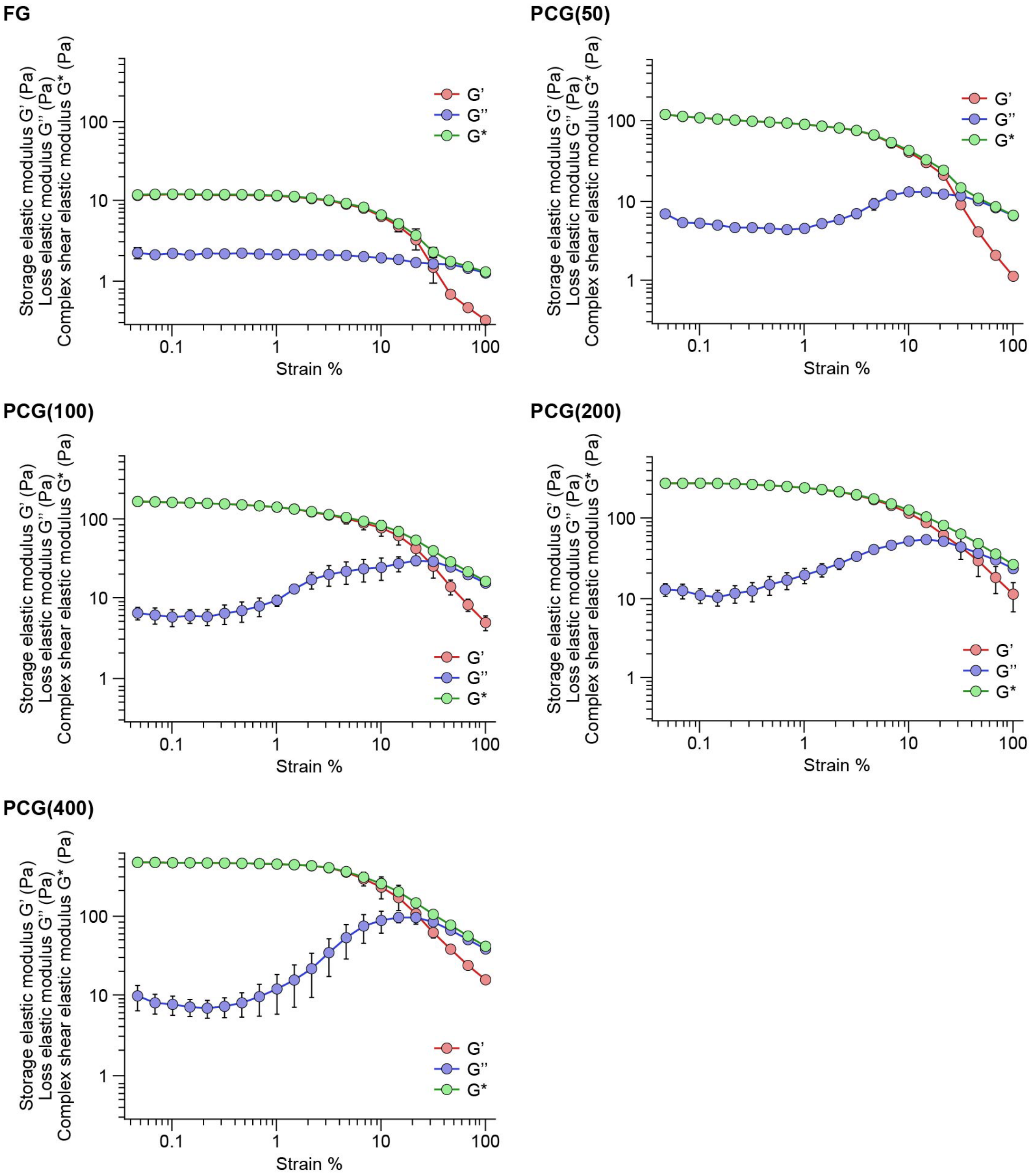


Figure S2. Dynamic viscoelastic characterization of collagen gels with varying TP(DMSO) concentrations. Dynamic viscoelastic properties of 2 mg/mL collagen gels containing different concentrations of TP(DMSO) were evaluated using a rheometer. Oscillatory shear measurements were performed at a fixed frequency of 1 Hz while the strain increased from 0.01% to 100%. Storage (G′), loss (G″), and complex shear modulus (G*) were recorded. Data represent mean ± standard error (n = 3).

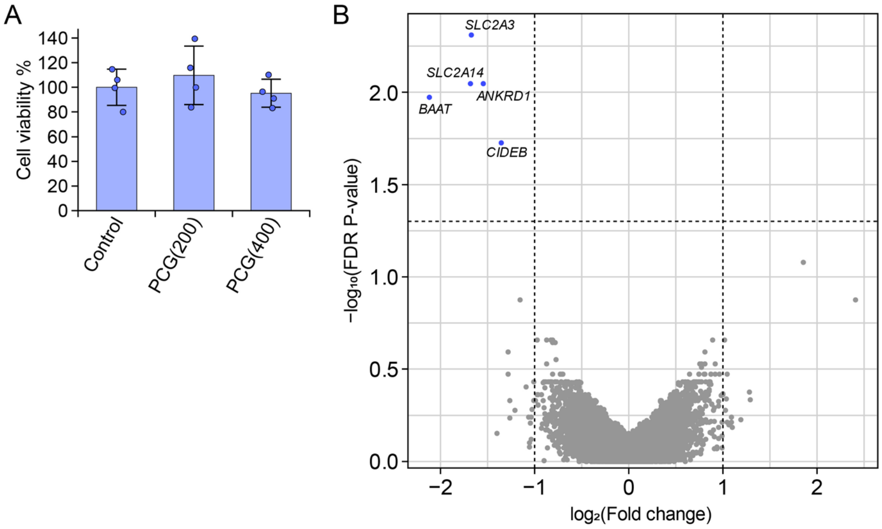


Figure S3. Evaluation of effects of platinum complexes eluted from PCGs. (A) Evaluation of cell viability. Free platinum complexes contained in PCGs were extracted into culture medium overnight. The extraction medium was then added to A549 cells cultured in a 96-well plate, followed by incubation for 1 day. Bars and error bars represent the mean ± standard deviation. (B) Gene expression analysis by DNA microarray. A549 cells were cultured in either standard culture medium or PCG(400) extraction medium for 1 day. A volcano plot was generated by comparing the PCG(400) extraction condition with the control condition following to DNA microarray analysis of isolated total RNA.

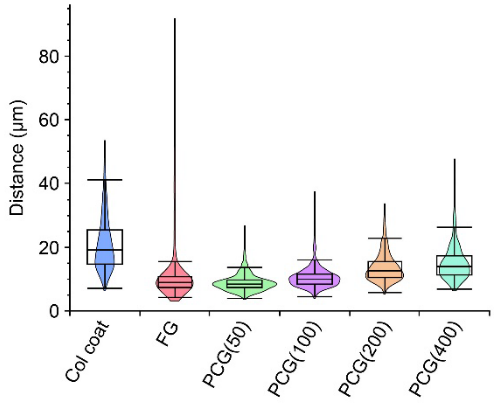


Figure S4. Nearest-neighbor distance profiles of A549 cells cultured on different substrates. Cells were cultured on the indicated substrates for 3 days, followed by staining nucleus with Hoechst 33342. Fluorescence images were acquired, and the coordinates of more than 700 nuclei per condition were measured. For each nucleus, the distance to the nearest neighboring nucleus was calculated. The distributions of the distances are presented as violin plots and box-and-whisker plots.

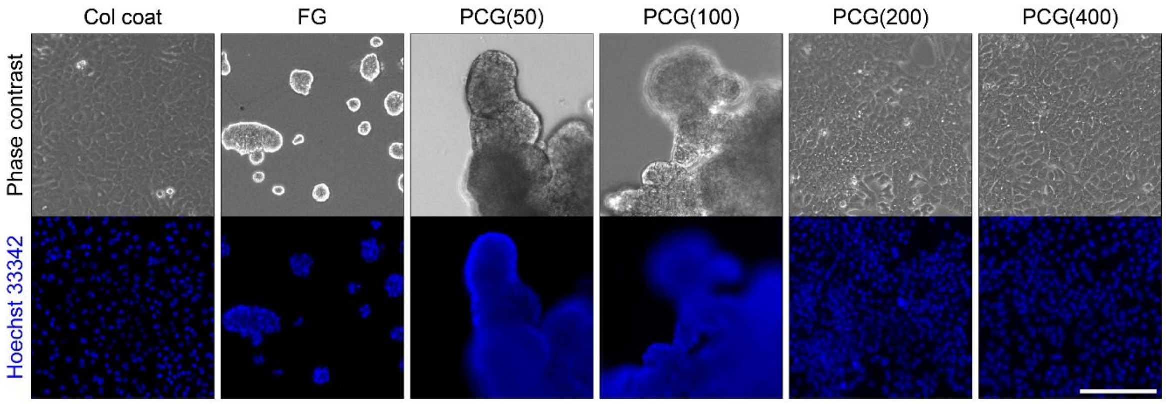


Figure S5. Representative fluorescence images of A549 cells on day 5 in the cell proliferation assay. Cells were cultured on the indicated substrates for 5 days for the cell proliferation assay shown in Figure 2C. Cell nuclei were stained with Hoechst 33342 and imaged using a fluorescence microscope. Scale bar represents 200 μm.

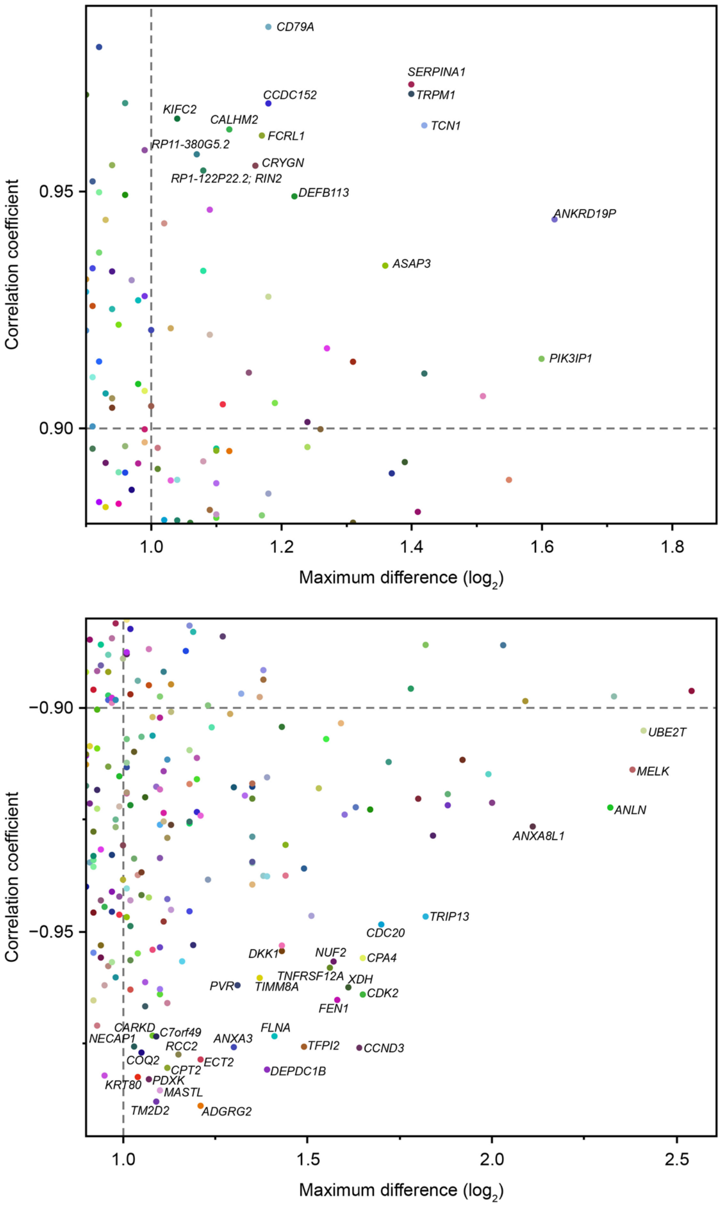


Figure S6. Correlation analysis between gene expression and cell aggregation. Gene expression data obtained from A549 cells cultured on different substrates were subjected to correlation analysis with the quantitative cell aggregation scores shown in Figure 2A. Genes showing positive or negative correlations are displayed in the upper and lower panels.

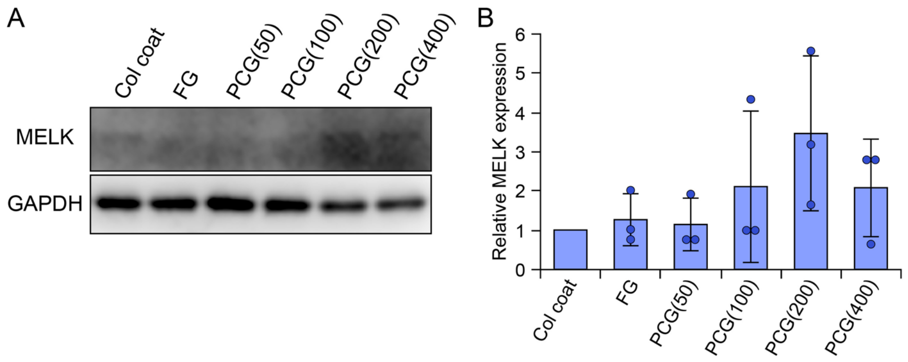


Figure S7. Western blotting of MELK in A549 cells cultured on different substrates. (A) Representative western blot images of MELK protein in A549 cells cultured on the indicated substrates for 2 days. (B) Densitometric analysis of MELK protein expression. Band intensities were quantified and normalized to GAPDH. Bars and error bars represent mean ± standard deviation (n = 3).

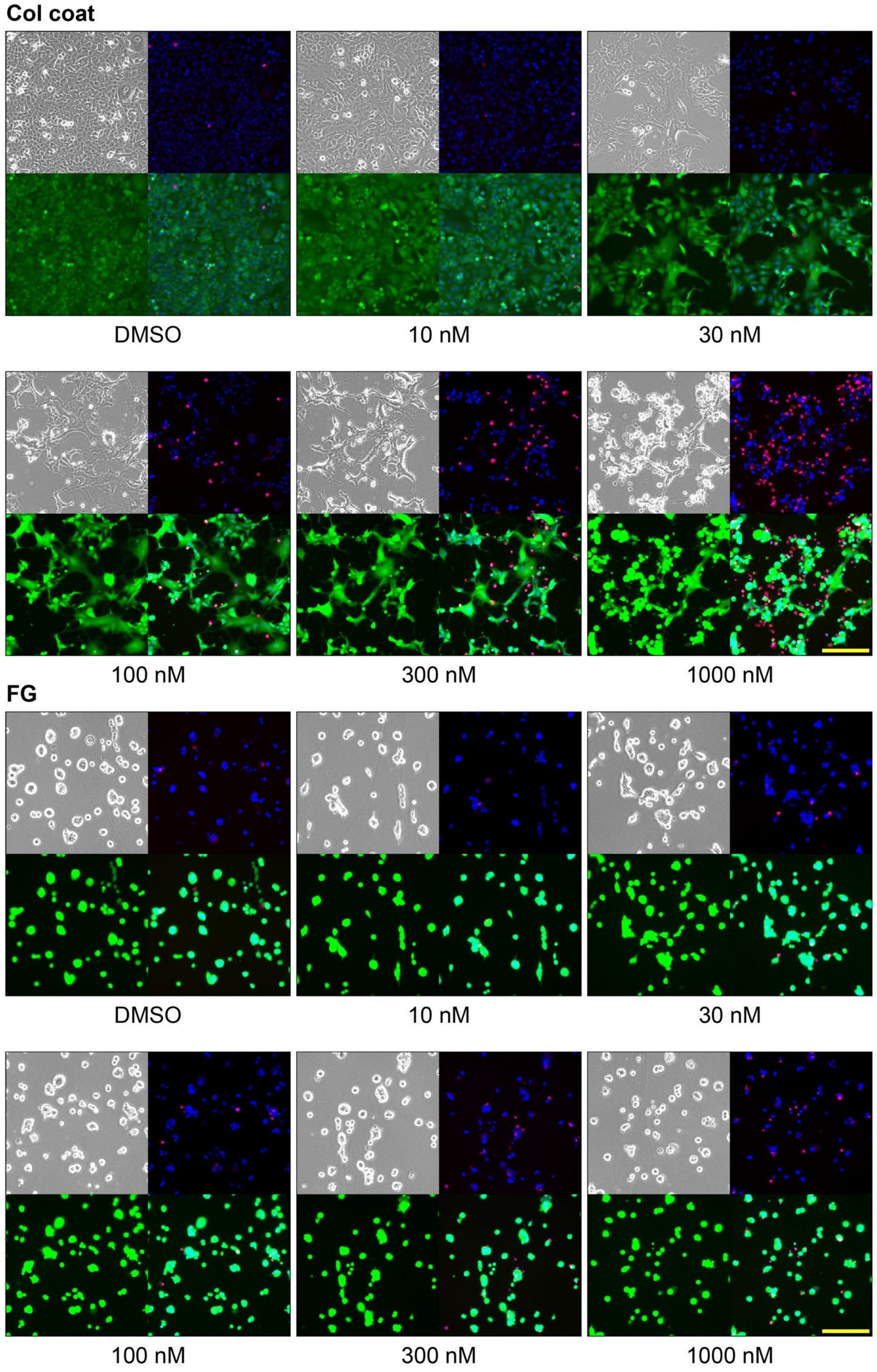


Figure S8. Fluorescence images of A549 cells after treatment with the MELK inhibitor OTSSP167 on different substrates. Cells were cultured on the indicated substrates for 2 days, followed by treatment with OTSSP167 at the indicated concentrations for one additional day. Cells were then stained with calcein-AM (green) to label live cells, PI (red) to label dead cells, and Hoechst 33342 (blue) to label total cell nuclei, and imaged using a fluorescence microscope. Phase contrast image (top left), merged PI and Hoechst

33342 image (top right), calcein-AM image (bottom left), and merged three-color fluorescence image (bottom right). Scale bars represent 200 µm.

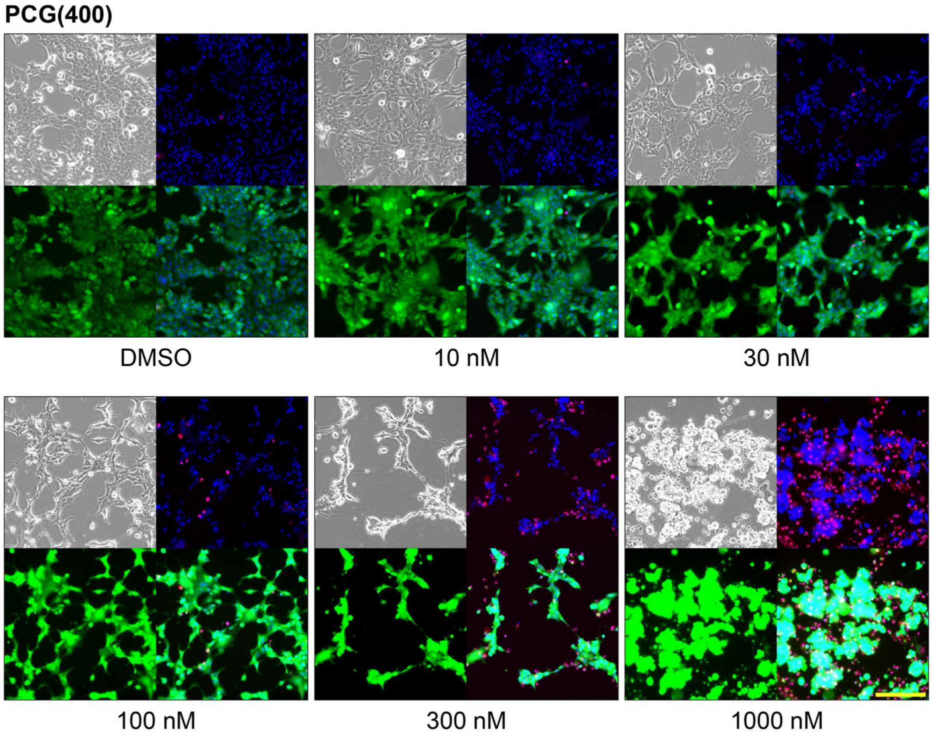


Figure S8. Continued

Table S1. List of tryptic peptides from bovine type I collagen α1 chain selected for quantification.

| Position | Sequence |
|---|---|
| −4–9 | ISVPGPMGPSGPR |
| 67–75 | GPOGPQGAR |
| 76–90 | GLOGTAGLOGMK[§]GHR |
| 109–126 | GEOGSOGENGAOGQMGPR |
| 133–144 | GROGAOGPAGAR |
| 145–174 | GNDGATGAAGPOGPTGPAGPOGFOGAVGAK[†] |
| 193–219 | GEOGPOGPAGAAGPAGNOGADGQOGAK |
| 220–237 | GANGAOGIAGAOGFOGAR |
| 238–252 | GPSGPQGPSGPOGPK[†] |
| 253–264 | GNSGEOGAOGSK |
| 295–309 | GEOGPAGLOGPOGER |
| 316–327 | GFOGADGVAGPK |
| 343–360 | GSOGEAGROGEAGLOGAK |
| 361–374 | GLTGSOGSOGPDGK |
| 375–386 | TGPOGPAGQDGR |
| 387–396 | OGPOGPOGAR |
| 397–408 | GQAGVMGFOGPK[†] |
| 421–434 | GVOGPOGAVGPAGK |
| 435–453 | DGEAGAQGPOGPAGPAGER |
| 508–519 | GVQGPOGPAGPR |
| 520–531 | GANGAOGNDGAK |
| 532–555 | GDAGAOGAOGSQGAOGLQGMOGER |
| 556–564 | GAAGLOGPK |
| 574–581 | GADGAOGK |
| 604–618 | GEAGPSGPAGPTGAR |
| 625–648 | GEOGPOGPAGFAGPOGADGQOGAK |
| 649–657 | GEOGDAGAK |
| 658–684 | GDAGPOGPAGPAGPOGPIGNVGAOGPK[†] |
| 658–687 | GDAGPOGPAGPAGPOGPIGNVGAOGPK[‡]GAR |
| 688–704 | GSAGPOGATGFOGAAGR |
| 705–725 | VGPOGPSGNAGPOGPOGPAGK |
| 741–756 | OGEVGPOGPOGPAGEK |
| 757–780 | GAOGADGPAGAOGTPGPQGIAGQR |
| 793–806 | GFOGLOGPSGEOGK |
| 807–816 | QGPSGASGER |
| 817–836 | GPOGPMGPOGLAGPOGESGR |
| 837–848 | EGAOGAEGSOGR |
| 889–906 | GETGPAGPAGPIGPVGAR |
| 907–915 | GPAGPQGPR |
| 934–963 | GFSGLQGPOGPOGSOGEQGPSGASGPAGPR |
| 964–974 | GPOGSAGSPGK |

[†] hydroxylysine, [‡] galactosyl-hydroxylysine, [§] glucosyl-galactosyl-hydroxylysine

*Residue numbering begins with the triple-helical region of the chains.

*4-Hydroxyproline is denoted by the single-letter code O.

*Met-containing peptides and a peptide containing both Met and His are shown in red and purple, respectively.

Table S2. List of tryptic peptides from bovine type I collagen α2 chain selected for quantification.

| Position | Sequence |
|---|---|
| −4–9 | GGGPGPMGLMGPR |
| 10–42 | GPOGASGAOGPQGFQGPOGEOGEOGQTGPAGAR |
| 135–144 | VGAOGPAGAR |
| 145–192 | GSDGSVGPVGPAGPIGSAGPOGFOGAOGPK††GELGPVGNOGPAGPAGPR |
| 193–219 | GEVGLOGLSGPVGPOGNOGANGLOGAK† |
| 238–252 | GIOGPVGAAGATGAR |
| 253–264 | GLVGEOGPAGSK |
| 292–309 | GSTGEIGPAGPOGPOGLR |
| 324–333 | AGVMGPAGSR |
| 343–360 | GPNGDSGROGEOGLMGPR |
| 361–374 | GFOGSOGNIGPAGK |
| 387–396 | OGPIGPAGAR |
| 397–408 | GEOGNIGFOGPK |
| 409–416 | GPSGDOGK |
| 421–429 | GHAGLAGAR |
| 430–453 | GAOGPDGNNGAQGPOGLQGVQGGK |
| 454–483 | GEQGPAGPOGFQGLOGPAGTAGEAGKOGER |
| 484–498 | GIOGEFGLOGOAGAR |
| 502–519 | GPOGESGAAGPTGPIGSR |
| 520–531 | GPSGPPGPDGNK |
| 556–564 | GAAGIOGGK |
| 586–603 | GAOGAIGAOGPAGANGDR |
| 604–618 | GEAGPAGPAGPAGPR |
| 625–648 | GEVGPAGPNGFAGPAGAAGQOGAK† |
| 658–687 | GENGPVGPTGPVGAAGPSGPNGPOGPAGSR |
| 688–704 | GDGGPOGATGFOGAAGR |
| 705–725 | TGOOGPSGISGPOGPOGPAGK |
| 741–756 | SGETGASGPOGFVGEK |
| 757–789 | GPSGEOGTAGPOGTOGPQGLLGAOGFLGLOGSR |
| 793–816 | GLOGVAGSVGEOGPLGIAGPOGAR |
| 817–836 | GPOGNVGNOGVNGAOGEAGR |
| 837–848 | DGNOGNDGPOGR |
| 859–884 | GYOGNAGPVGAAGAOGPQGPVGPVGK |
| 889–906 | GEOGPAGAVGPAGAVGPR |
| 934–963 | GHNGLQGLOGLAGHHGDQGAOGAVGPAGPR |
| 964–974 | GPAGPSGPAGK |
| 978–990 | IGQOGAVGPAGIR |

†hydroxylysine,
††galactosyl-hydroxylysine/glucosyl-galactosyl-hydroxylysine (averaged)
*Residue numbering begins with the triple-helical region of the chains.
*4-Hydroxyproline is denoted by the single-letter code O.
*Met-containing peptides and His-containing peptides are shown in red and blue, respectively.

Table S3. Gene lists highly correlated with compressive elastic modulus.

| Correlated with | Gene | Correlation coefficient | Description |
|---|---|---|---|
| Compressive modulus | *ID3* | 0.992 | Negative regulation of transcription |
| | *ACIN1* | 0.992 | Apoptosis, RNA processing |
| | *POLR3B* | 0.988 | RNA polymerase |
| | *HIST1H2BM* | 0.987 | Nucleosome |
| | *ESCO1* | 0.985 | Cell cycle |
| | *ABCC5* | 0.985 | ABC transporter |
| | *ZW10* | 0.984 | Cell cycle |
| | *HIST1H2AG* | 0.983 | Nucleosome |
| | *PTCD2* | 0.982 | Cell surface interactions |
| | *MAP3K4* | 0.981 | MAPK cascade |
| | *CEP89* | 0.980 | Centrosome |
| | *CXXC5* | 0.980 | Estrogen receptor signaling |
| | *IGFL3* | 0.978 | IGF-like |
| | *INSL4* | 0.978 | Insulin-like |
| | *ARPC5L* | 0.978 | Actin polymerization |
| | *BAD* | 0.977 | Apoptosis |
| | *HIST1H2AK* | 0.976 | Nucleosome |
| | *PLCZ1* | 0.975 | Ca2+ release |
| | *GINS2* | 0.974 | DNA replication |
| | *LY96* | 0.973 | Immune response |
| | *CAPN9* | -0.997 | $Ca^{2+}$-dependent proteolysis |
| | *ZNF660* | -0.995 | Transcription factor |
| | *STRA8* | -0.992 | Splicing |
| | *LRG1* | -0.992 | Signal transduction |
| | *SERPINE3* | -0.988 | Serine protease inhibitor |
| | *CPB2* | -0.985 | Peptidase |
| | *DDR1* | -0.984 | Collagen receptor tyrosine kinase |
| | *TMEM240* | -0.983 | Unknown |
| | *OR8D2* | -0.983 | Olfactory receptor |
| | *NLRP9* | -0.982 | Inflammasome activation |
| | *DSPP* | -0.980 | Dentinogenesis |
| | *RNASE6* | -0.980 | RNA nuclease |
| | *MFSD2B* | -0.977 | Sphingolipid transporter |
| | *IL20RB* | -0.976 | IL-20 receptor |
| | *PLEKHG4B* | -0.974 | RhoGEF |
| | *PSMB2* | -0.974 | Proteasome |
| | *FGD1* | -0.973 | RhoGEF |
| | *POMC* | -0.972 | Proopiomelanocortin |
| | *PAX2* | -0.971 | Transcription factor |
| | *ARHGAP15* | -0.969 | RhoGAP |

Table S4. Gene lists highly correlated with shear elastic modulus.

| Correlated with | Gene | Correlation coefficient | Description |
|---|---|---|---|
| Shear modulus | *TM7SF3* | 0.998 | Negative regulation of apoptosis |
| | *PFKP* | 0.996 | Glycolysis |
| | *CSTF2* | 0.995 | RNA processing |
| | *OR51E1* | 0.993 | Olfactory receptor |
| | *FAM136A* | 0.993 | Respiration electron transport |
| | *ATP6V1G2-DDX39B* | 0.992 | ATPase proton transporter |
| | *DUT* | 0.992 | Nucleic acid metabolism |
| | *PDHA1* | 0.992 | Glycolysis |
| | *FBXW10* | 0.991 | Ubiquitynation |
| | *FEN1* | 0.990 | DNA repair, DNA replication |
| | *CXCL6* | 0.989 | Chemokine |
| | *TRIM16L* | 0.989 | Pseudogene |
| | *PTBP2* | 0.989 | RNA processing |
| | *SEMA3A* | 0.988 | Axon guidance |
| | *SLC16A1* | 0.988 | Monocarboxylate transporter |
| | *TIMM8A* | 0.987 | Translocase |
| | *GLDC* | 0.987 | Glycine metabolism |
| | *CENPF* | 0.987 | Kinetochore |
| | *NIPSNAP1* | 0.986 | Mitochondria homeostasis |
| | *KCND1* | 0.986 | Potassium channel |
| | *UNC79* | -0.998 | $Na^+$ channel |
| | *PRPF40B* | -0.997 | RNA processing |
| | *DEFA3* | -0.996 | Defense response |
| | *SERPINA1* | -0.995 | Anti-inflammation |
| | *ACAP3* | -0.995 | ArfGAP |
| | *TBC1D30* | -0.988 | RabGAP |
| | *VSTM2A* | -0.988 | Positive regulation of transcription |
| | *A2BP1* | -0.985 | RNA processing |
| | *KIFC2* | -0.985 | Motor protein |
| | *SCARA5* | -0.985 | Ferritin receptor |
| | *NLRP10* | -0.985 | Inflammasome |
| | *ANKRD19P* | -0.982 | Pseudogene |
| | *ASAP3* | -0.981 | ArfGAP |
| | *TCN1* | -0.980 | Cobalamin transport |
| | *SLC25A19* | -0.977 | Thiamine pyrophosphate transport |
| | *PRDM6* | -0.975 | Histone methylation |
| | *ANGPTL3* | -0.975 | Metabolic hormone |
| | *RIN2* | -0.974 | RabGEF |
| | *TP63* | -0.974 | Transcription factor |
| | *OR10A5* | -0.974 | Olfactory receptor |
